# Parameterisation of epidemiological models from small field experiments: a case study of banana bunchy top virus transmission

**DOI:** 10.1101/2025.11.11.687876

**Authors:** Renata Retkute, Aman Bonaventure Omondi, Misheck Soko, Charles Staver, John E. Thomas, Christopher A. Gilligan

## Abstract

Accurate estimation of epidemiological parameters from limited field data remains a major challenge in plant disease modeling. We present a novel data-augmented adaptive multiple importance sampling (DA-AMIS) framework that integrates Bayesian inference with stochastic epidemic modeling to estimate key transmission parameters from small field experiments. Using detailed individual-level observations from a 24-plant experiment on the natural spread of banana bunchy top virus (BBTV) in Benin, we jointly inferred infection timing, dispersal characteristics, and transmission rates for both primary and secondary infections. Model validation against independent datasets from BBTV field trials in Burundi and Malawi showed close correspondence between simulated and observed prevalence dynamics, confirming the generality of parameter estimates across regions. The inferred 12% infection rate of replanting suckers underscores the risk of disease introduction through planting material, while simulations identified April as the period of peak infection, providing actionable insights for surveillance timing.

## 1 Introduction

Mathematical models of epidemics can provide insights into the mechanisms underlying the spread of disease and allow comparison of different management scenarios (Anderson and May, 1991). Significant challenges remain, however, in the estimation of parameters of epidemiological models to inform disease management (Swallow et al., 2022). This is especially true for forecasting of emerging and reemerging plant diseases, since epidemiological data are usually incomplete (Cunniffe et al., 2014). Typically, these data consist of successive surveys at fixed time periods, in which the disease status is assessed on subsamples of susceptible hosts within a target population. As such, the exact times of infection are never observed in contrast with epidemics in human populations where data are frequently recorded for onset of symptom expression (Cauchemez and Ferguson, 2008).

The scarcity and coarse temporal resolution of plant disease data present a fundamental limitation for quantitative inference. When infection events cannot be directly observed, estimates of key transmission parameters—such as infection rate, latent period, and dispersal—must rely on indirect or aggregated observations, often leading to considerable uncertainty in model fitting and prediction (Parnell et al., 2012). Small field experiments could offer a practical and efficient approach for estimating transmission parameters under natural conditions. Such experiments are comparatively inexpensive to establish and manage, require fewer resources for disease surveillance than extensive landscape-scale surveys, and can yield higher-quality data with reduced operator error by allowing detailed assessment of individual plants. However, despite these potential advantages, their use for parameter estimation has rarely been explored formally, and few statistical frameworks have been developed to address model parameterisation from limited or censored epidemiological data.

Banana bunchy top disease (BBTD) is the most damaging viral disease of banana worldwide that, in particular, poses one of the greatest threats to banana cultivation by smallholder farmers (Kumar et al., 2011). Currently, there are no banana cultivars or landraces fully resistant to BBTV (Rahayuniatia and Subandiyahb, 2022; Cueva et al., 2023). Infection can cause up to 100% yield loss (Okonya et al., 2019),, which may lead to the loss of seed businesses due to infection risk labeling and the disappearance of farmer cultivars (Simbare et al., 2020). The virus is transmitted from infected to healthy plants by an aphid vector *Pentalonia nigronervosa* (Magee, 1927, 1940) with additional transmission to field sites by the use of infected propagation material (Kumar et al., 2011). The virus has spread rapidly in sub-Saharan Africa during the past two decades Ploetz et al. (2015). Large areas covering mainly Central Africa, parts of West Africa, and areas within East Africa are at high risk of the establishment and subsequent spread of BBTV Bouwmeester et al. (2023).

In this paper, we investigate how to make optimal use of small field studies to parameterise epidemic transmission models. We develop a new approach that combines adaptive multiple importance sampling (AMIS) (Cornuet et al., 2012) with data augmentation methods (Jewell et al., 2008; Adrakey et al., 2017; Prangle and Viscardi, 2023). Our methodology evaluates uncertainty in the estimated parameters by assessing the likelihood of the empirical data based on sampled parameters. Adaptive multiple importance sampling has been used for parameter estimation in a variety of research fields, including epidemiology (Retkute et al., 2021; Abboud et al., 2023), physics (Prusokiene et al., 2021; Llorente et al., 2024), biology (Burgess et al., 2023; Prusokiene et al., 2023) and environmental sciences (Retkute et al., 2024). To parameterise virus spread in a discrete population, we use individual-level data from a small field experiment, with the data augmentation framework additionally enabling inference on unobserved host properties, such as infection timing.

For parameterisation, we utilize data from field experiments on the natural spread of BBTV among small populations of banana plants and validate the model using independent data from a larger-scale experiment. Our analyses identify several factors influencing the implementation of BBTD control strategies. Specifically, we estimate the probability that suckers sourced from the field are infected, highlighting the risk of virus introduction through planting material. We also determine the periods when the expected number of new infections is highest. These findings offer a practical framework for deriving epidemiological parameters from field experiments and can help guide more effective disease management strategies, ultimately supporting sustainable banana production in areas affected by BBTD.

## 2 Methods

### 2.1 Experiment used for model parameterisation

We used data for model parameterisation from an experiment that was conducted at IITA Bénin experimental gardens (lat=6^°^25.15 N, lon=2^°^19.67 E). The field was established on 1st December 2016. The plants were arranged in a rectangular frame, with an inter-row distance of 3 m and a between-column distance of 2 m. The experiment relied on natural infection from surrounding endemically infected plantations. Surveillance was conducted each month from June 2018 to June 2021. When BBTV was detected, mats were removed and plants replaced by suckers (offshoots, that grow from the base of the banana plant) from the same field,, which appeared visually healthy. The date and location (row and column IDs) of all removed banana plants were recorded throughout the duration of the experiment. Therefore for each *i* = 1 : *N* plant we have *x* and *y* coordinates, date of planting and date of removal. We will call this the Benin dataset.

### 2.2 Data for model validation

For model validation, we used two published datasets: (i) an experiment conducted in Burundi (Niyongere et al., 2012), and (ii) an experiment conducted in Malawi (Omondi et al., 2020). The Burundi experiments were carried out on small-scale banana farms and included multiple cultivars. Each plot measured 15 m ×8 m with inter- and intra-row plant spacing of 3 m and 2 m. The incidence of BBTD was monitored over nine months. We will call this the Burundi dataset. The Malawi experiments spanned three years, with banana plants established at 3 m between rows and 2.5 m between columns. These were natural infection experiments, with infections arising from external sources, and field-level BBTD incidence (%) was recorded during monthly surveys. We will call this the Malawi dataset.

### 2.3 BBTV transmission model

We model the spread of BBTV using a spatio-temporal, individual-based stochastic model. Each plant is classified in one of five non-overlapping compartments: susceptible (S), exposed (E), infectious (I), detected (D) and removed (R). A model schematic is shown in Figure 1. The model can account for instant removal after detection or delayed removal, for example when there are logistical constraints following inspection. Removed plants are replanted using suckers sourced from the field or externally, which may be healthy or infected.

**Figure 1.**
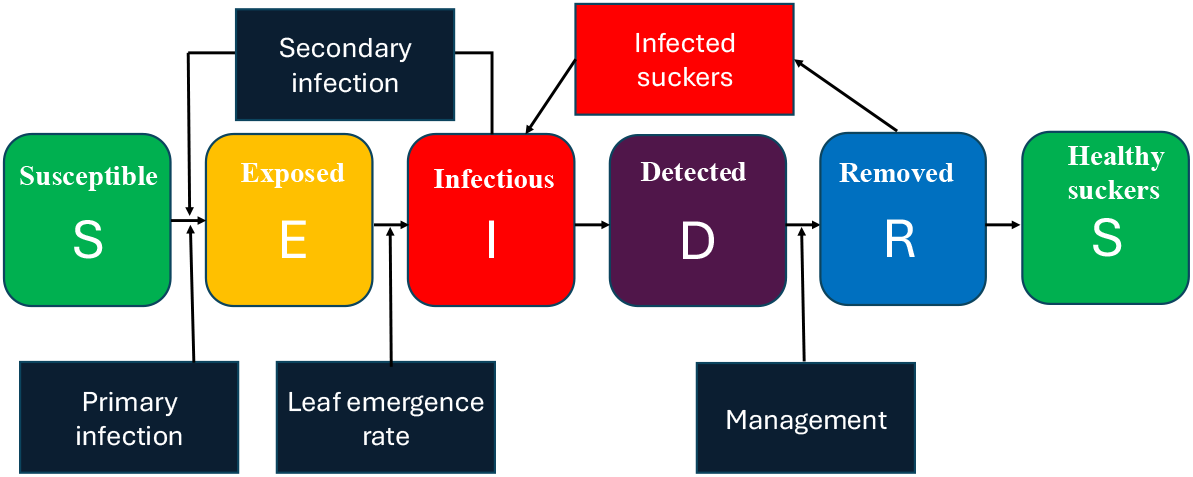
Schematic diagram for the BBTV transmission model: Susceptible (S), Exposed (E), Infectious (I), Detected (D), and Removed (R), with three sources of infection: primary infection originates from external inoculum sources, infected suckers for re-planting and secondary infection by insect vector transmission.

The spread of BBTV involves two processes: primary infection, in which the virus is introduced from outside the plantation area, and secondary infection between infected and healthy plants within the plantation area. The force of infection on a susceptible plant *i* at time *t* is given by

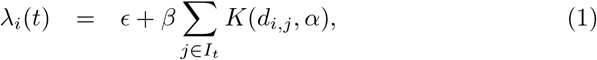

where *ϵ* is the primary infection rate, *β* is the secondary infection rate, *K*(*d, α*) is the dispersal kernel for secondary infection, *d*_*i,j*_ is the distance between plant *i* and plant *j*, and *I*_*t*_, is an index set of infectious plants at time *t*. We define the dispersal kernel as:

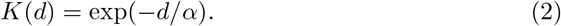

The latent period of BBTV is equal to the time required to produce 3.7 banana leaves (Allen, 1987). These periods can be inferred if leaf emergence rate (LER) is known. The leaf emergence rate has seasonal dependence and can be expressed as a function of day of the year, following (Allen et al., 1988):

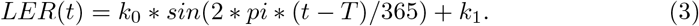

We used values for the parameters, *k*_0_ = 0.056, *k*_1_ = 0.062, from (Allen et al., 1988), while setting *T* = 90 days to align LER with the onset of the rainy season in West Africa (Laux et al., 2008).

We assumed that *p*% of suckers produced by infected but not yet infectious plants are infectious, and that 100% of suckers produced by infectious plants are infectious. Suckers are not tested for BBTV before planting, therefore representing an additional source of infection. We further assume that all infected plants eventually express symptoms and will ultimately be detected and removed, with removal following an exponential distribution with rate *γ day*^−1^. The model is simulated using the tau-leaping algorithm (Gillespie, 2001).

### 2.4 Complete-data likelihood

Let *t*_obs_ denote the duration of the observation period. We label the hosts that became infected by *i* = 1, 2, …, *n*_*I*_ and the remaining uninfected hosts by *i* = *n*_*I*_ + 1, *n*_*I*_ + 2, …, *N*. The planting time of plant *i* is denoted by 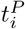. For infected plants, we denote by 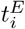 the time of infection, 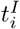 the time the plant becomes infectious, and 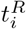the time of removal. We assume that the infectious period duration follows an exponential distribution, 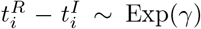, with removal rate *γ*.

At any time *t*, we define the sets of susceptible, exposed and infectious plants as:

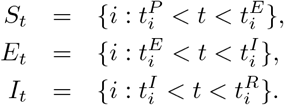

Assuming complete data Ω = {*S*_*t*_, *E*_*t*_, *I*_*t*_} for all *t* ∈ [0, *t*_obs_] and model parameters *θ* = {*α, β*,, *γ, p*}, we can express the likelihood as:

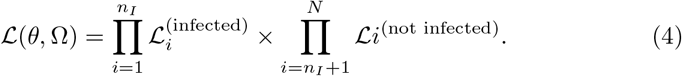

The components of the Eq.(4) are calculated as follows.

The cumulative infection pressure experienced by plant *i* from planting time 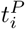 up to its infection time 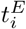 is calculated as:

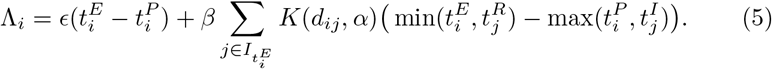

For plant *i*, which was planted at the start of the experiment, the likelihood that it became infected at time 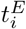 and was removed at time *t*_*i*_*R* is equal:

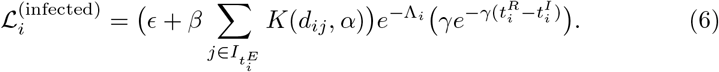

For planted suckers, they may have been infected at the time of planting (with probability *p*), or they may have been healthy initially but became infected later through primary or secondary infection. The likelihood of infection and removal is therefore expressed as:

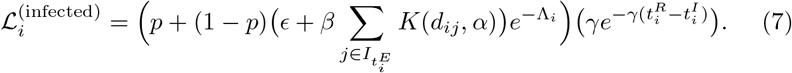

Likelihood for plant *i*, which was not infected during time from 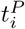 to *t*_*obs*_ is:

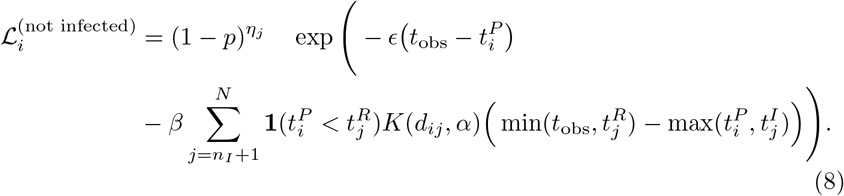

The indicator *η*_*i*_ = 1 if plant *i* was planted as a sucker during the experiment, and **1**(*x*) = 1, if condition *x* is satisfied, and zero otherwise.

### 2.5 Data augmented adaptive multiple importance sampling (DA-AMIS)

For parameter estimation, we developed a novel approach that combines adaptive multiple importance sampling (AMIS) (Cornuet et al., 2012) with data augmentation methods (Jewell et al., 2008; Adrakey et al., 2017). Using a proposal constructed from an AMIS algorithm, we sample parameters { *α, β*,, *γ, p}*. Next, based on the value of parameter *γ*, we propose a date when plants became infectious, *t*^*I*^, and back-calculate the time of infection, *t*^*E*^, using values of leaf emergence rate prior to time *t*^*I*^. Finally, we calculate the value of the likelihood function given by Eq.4.

For each iteration *l* = 1, …, *L*, we sample *N*_*l*_ parameter sets. At the first iteration, we sample parameters from the prior distribution. We chose uniform priors for each of the five model parameters (Table 1). At iterations *l* = 2, …, *L*, a suitable proposal is found by fitting a mixture of Student’s t-distributions to the weighted samples. The corresponding importance weights for a sampled set of parameters *θ*_*k*_ at iteration *l* can be calculated as (Cornuet et al., 2012):

**Table 1:** Definitions and prior distributions of model parameters.

| Parameter | Definition | Prior distribution |
| --- | --- | --- |
| $\epsilon$ | Primary infection rate ($day^{-1}$) | $U[0, 1]$ |
| $\beta$ | Secondary infection rate ($day^{-1}$) | $U[0, 1]$ |
| $\alpha$ | Dispersal kernel parameter ($m$) | $U[0, 25]$ |
| $\gamma$ | Removal rate ($day^{-1}$) | $U[1/100, 1/10]$ |
| $p$ | Proportion of infected suckers | $U[0, 1]$ |

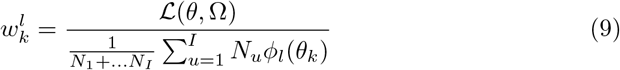

where *φ*_*l*_(*θ*_*k*_) is the proposal distribution at iteration *l*. We use Kish’s effective sample size (ESS) (Kish, 1966) as a measure of the quality of the posterior distribution:

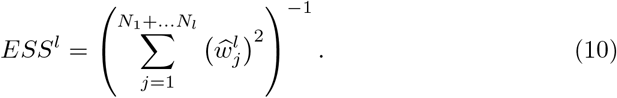

The algorithm continues until the *ESS* is above the required threshold, or the maximum number of iterations, *L*, has been completed.

### 2.6 Model accuracy assessment

The probabilistic model accuracy was evaluated using the weighted interval score (WIS), a non-negative metric, which measures how consistent predicted distributions are with observed values (Bracher et al., 2021). Given an observation *y*, an uncertainty level *a*, and a simulated distribution *F*, an interval score is defined as

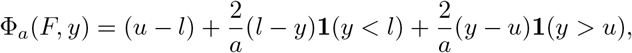

*l* and *u* are the 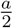 and 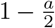 quantiles of simulated distribution *F*. The interval score has a low value when the simulated distribution is not-dispersed (l and u are close), but penalises cases when the observed value lies outside the interval [*l, u*]. The interval scores corresponding to several uncertainties can be combined into a single metric, i.e. weighted interval score (WIS). Then WIS is defined as a linear combination of *K* interval scores:

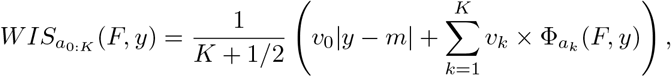

where weights 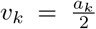 for *k* = 1, …, *K* and *v*_0_ = 1*/*2, and *m* is the median of the simulated distribution. We used *K* = 11 interval scores with *a* = 0.02, 0.05, 0.1, …, 0.9 (Bracher et al., 2021). Model simulations were performed using 1,000 parameter sets randomly drawn from the posterior distribution.

## 3 Results

### 3.1 Small field experiment on BBTV transmission

In the course of the experiment, 16 plants were found to be infected, based upon symptom expression. The number of infected plants reported at monthly surveys and the locations of susceptible and infected plants are shown in Fig.2. During most survey dates either no infected plants were found, or only a single infected plant was detected, with the exception of June 2018 (4 infected plants), November 2018 (2 infected plants) and February 2020 (2 infected plants).

**Figure 2.**
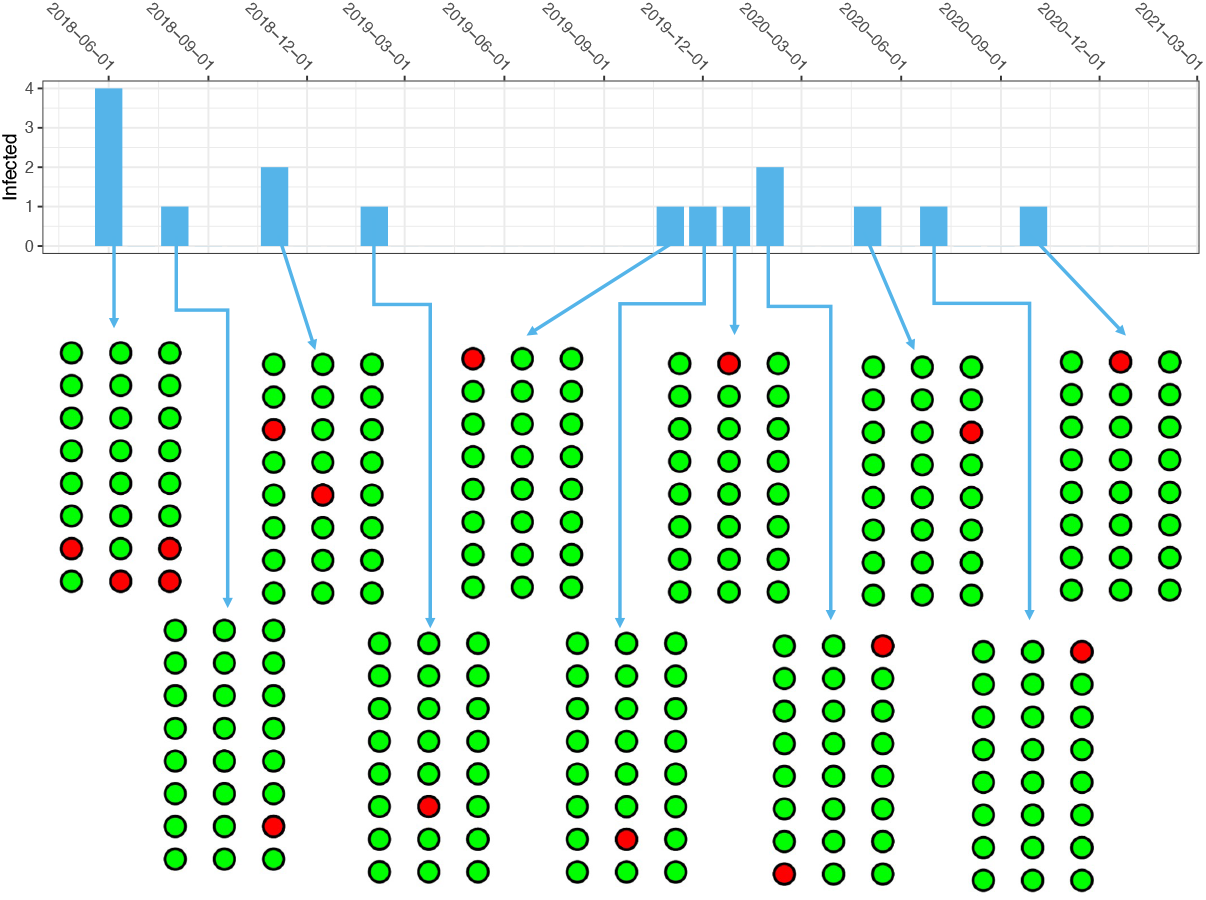
Experimental data: number of infected plants reported at monthly surveys and locations of susceptible (green circles) and infected (red circles) plants. Distance between columns is 3 meters and between rows in 2 meters.

### 3.2 Parameterisation of the BBTV transmission model

We set the number of parameters sampled to *N*_*l*_ = 1, 000 for *l* = 1, …, *L*. The maximum number of iterations was set to *L* = 1, 000, and the required effective sample size to *ESS*_*R*_ = 1, 000. Uniform priors were chosen for the model parameters (Table 1). Calculations were run on a single MacBook Pro with a 2.4 GHz 8-Core Intel Core i9 processor. The number of iterations required to achieve *ESS* ≥1, 000 was 25, which took approximately 5 minutes.

The performance of the DA-AMIS algorithm was assessed by examining the evolution of likelihood values and the ESS across iterations. The distribution of likelihood values (Figure 3A) shows that, over successive iterations, the algorithm increasingly concentrates on high-likelihood regions of the parameter space. During the first 10 iterations, the range of log-likelihood values was (−48, 110; −218), reflecting the wide range of the prior distribution. After iteration 10, refinement of the posterior exploration progressively reduced the log-likelihood range to (−211; −179). Correspondingly, the ESS (Figure 3B) increased over iterations, reflecting improvements in sampling efficiency and convergence of the algorithm.

**Figure 3.**
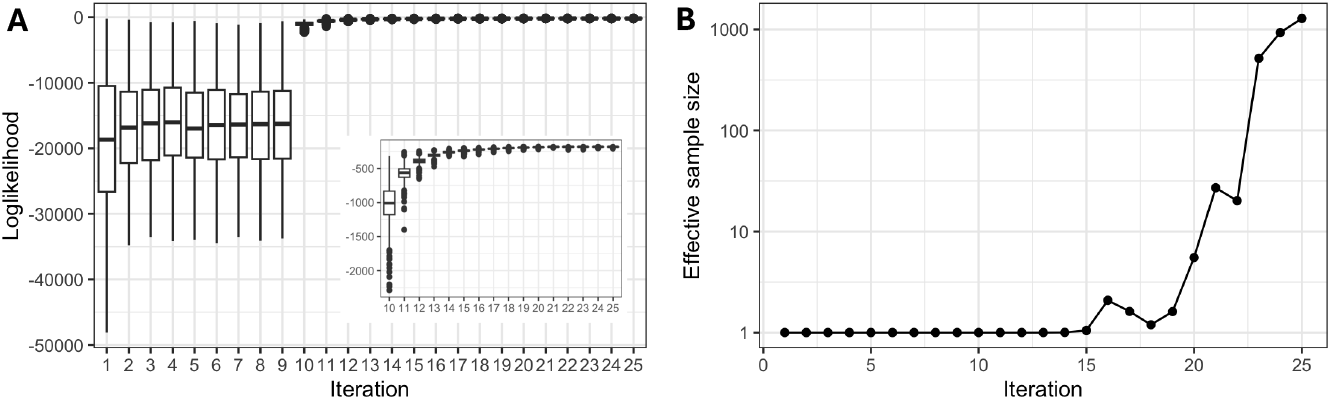
Performance of DA-AMIS algorithm. (A) Distribution of likelihood values as a function of iteration. Insert shows loglikelihood distributions for iterations 10 − 25. (B) Change in ESS as a function of iteration.

The posterior distributions of the model parameters are shown in Figure 4. The mean values for the primary and secondary transmission rates were estimated to be 4.3 ×10^−4^ per day, and 6.8 ×10^−4^ per day, respectively. The marginal distribution of the primary infection rate had a nicely defined unimodal shape. The marginal distribution of the secondary infection rate did not have a well defined peak; however, our results provide what could be an expected upper bound for the rate of secondary infection. The mean dispersal kernel parameter had a value 14.6m. The mean proportion of planted suckers that were infected was 0.12 (95% CI 0.03-0.25).

**Figure 4.**
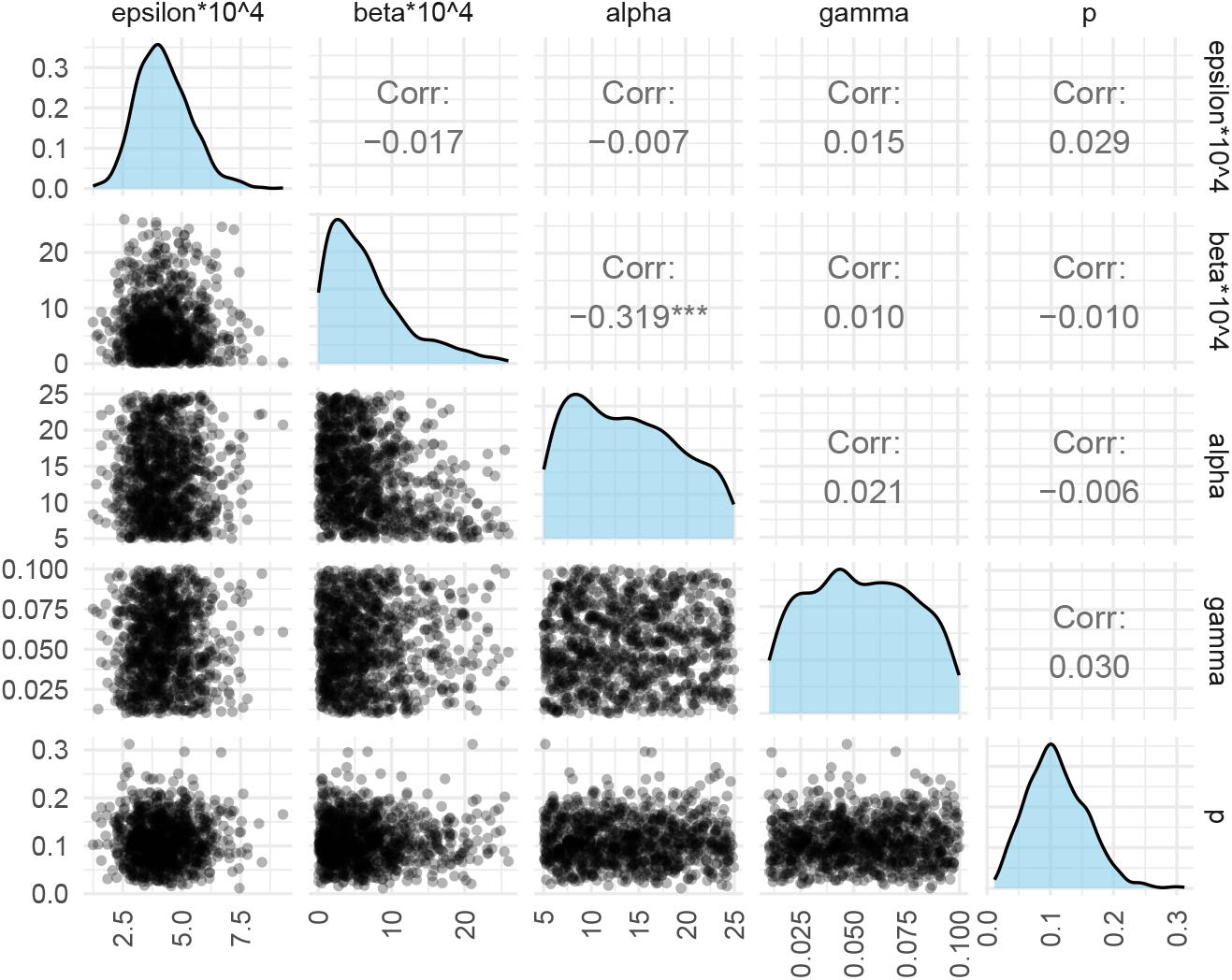
The posterior distributions of the model parameters: primary infection rate (ϵ), secondary infection rate (*β*), dispersal kernel parameter (*α*), removal rate (*γ*), and proportion of infected suckers (*p*). See Table 1 for priors.

### 3.3 Validation of the BBTV transmission model

Experiments conducted in Burundi involved 12 banana plants per plot (Niyongere et al., 2012), which was fewer than in the Benin experiment used for parameterisation of the epidemiological model. The incidence of BBTD in the Burundi experiments varied significantly with trial location, banana cultivar, and planting material type, with prevalence ranging from 0.7% to 56.4% nine months after establishment (Niyongere et al., 2012). The simulations showed good agreement with the experimental data (Fig. 5A) and captured the observed variation in prevalence throughout the experiment, with all observed values falling within the 95% confidence interval of the simulations.

**Figure 5.**
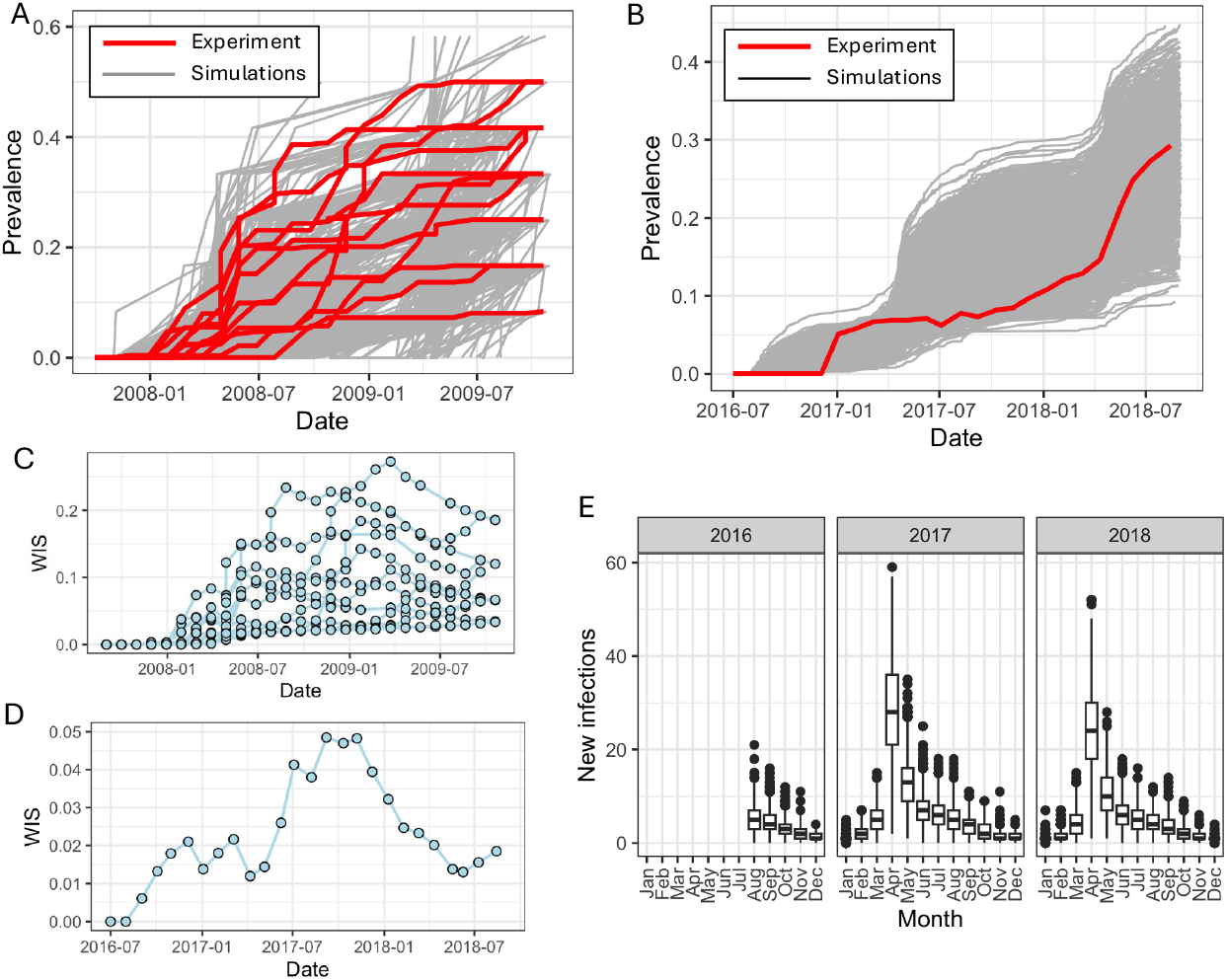
Comparing the field experiment data with model simulations. (A) Model simulations and experiment data for Burundi (Niyongere et al., 2012). (B) Model simulations and experiment data for Malawi (Omondi et al., 2020). The WIS as a function of date for simulated outbreaks for Burundi data. (D) The WIS as a function of date for simulated outbreaks for Malawi. (E) Boxplot distributions of predicted new infections as a function of year and month.

To compare disease dynamics with experimental data in the Malawi study (Omondi et al., 2020), epidemics were simulated on a representative field with 30 rows by 20 columns. This corresponds to a plot size of 0.45 ha, consistent with banana field sizes in Africa, where plots can range from 0.01 ha to 1 ha, with an average of 0.2 ha (Chabi et al., 2018). The simulations exhibited a wider range of prevalence, likely reflecting variation in the posterior distribution of the secondary infection rate (Fig. 5B). Overall, the model captured the field-observed dynamics of BBTV well, showing strong agreement between simulated and reported disease patterns.

To assess model performance as a function of time, we calculated the weighted interval scores at all survey dates. The WIS is a measure of how close the distribution from model simulation is to the observation (Bracher et al., 2021). Values of WIS closer to zero indicate a better performance of the model. For the Burundi data, 68% of WIS values were below 0.2 (Fig.5 C). The model showed a good performance (WIS*<*0.2) for Malawi data as well, with WIS values below 0.05 (Fig.5 D).

We used simulations based on the Malawi experiment configuration to track the number of new infections by month and year. This temporal resolution allowed us to identify periods optimal for BBTV detection during surveillance. Most new infections were predicted to occur in April (Fig. 5E), a pattern consistent across 2017 and 2018. The model’s predicted increase in BBTV prevalence between April and September 2018 aligned closely with the observed rise in prevalence from the experimental data.

## 4 Discussion

In the current study, we explored how small field experiments can be used to estimate epidemiological parameters for virus transmission and dispersal. In our experiment, the host population consisted of only 24 banana plants. To date, methods for estimating epidemiological parameters from such small-scale experiments have been limited, typically restricted to observational analyses of disease progress or fitting simple growth curves, such as logistic models (Berger, 1981; Gilligan, 1990).

Previous work on modeling the spread of banana bunchy top disease (BBTD) is limited (Allen, 1977, 1987; Smith et al., 1998; Varghese et al., 2020; Retkute and Gilligan, 2025; Mapinda et al., 2025c,b,a; Retkute et al., 2025). The study most closely related to ours is Varghese et al. (2020), which developed a stochastic network model of BBTV transmission using a large dataset from a plantation of approximately 24,000 plants, of which over 3,000 were removed over three years. Parameter estimation in that study relied on approximate Bayesian computation (ABC), requiring 10^7^ parameter samples to achieve an acceptance rate of 0.004.

We have introduced a novel parameter estimation approach, DA-AMIS, which simultaneously infers epidemiological parameters and infection times. Using this method, we achieved an effective sample size of 1,000 with only ∼3 × 10^5^ parameter samples. Our estimates of the primary infection rate ranged from 1.3× 10^−4^ to 8.9 ×10^−4^ *day*^−1^, while the secondary infection rate provided an upper bound of 27.5× 10^−4^ *day*^−1^. The mean dispersal kernel was estimated at 14.6 m, closely matching the 17.2 m reported by Allen (1978). Replacement of removed plants with suckers from the same field allowed us to estimate that on average 12% of planting material was already infected, a critical parameter when modeling control strategies involving clean planting material.

We validated our model using independent data from both smaller and larger field experiment, observing strong agreement between simulated and fieldreported BBTV dynamics. Model performance, quantified using the weighted interval score, showed that simulated prevalence distributions closely matched observed values. Despite being parameterised from the small field experiment, the model captured key epidemic features, including changes in prevalence slope across epidemic phases. Some discrepancies were observed during the early epidemic, likely due to our assumption of 100% detection probability, whereas early symptomatic infections may have been missed in the field. Our simulations consistently identified April as the period with the highest expected number of new infections, corresponding to the peak and end of the rainy season in Malawi, which typically spans from November to April (Ngongondo et al., 2011). This predicted increase in BBTV prevalence likely reflects the favourable environmental conditions at this time of year, when the onset of rains and rising temperatures promote banana growth - producing more young, susceptible tissue—and enhance the activity and transmission efficiency of the banana aphid vector (Anhalt and Almeida, 2008). One other striking result was the ability to extend inferences from a field experiment in Benin in West Africa to model dynamics in fields experiments in Burundi and Malawi in East Africa. This represents a considerable extrapolation, highlighting the challenges and importance of scaling up from localized experiments. Data that record disease progress in carefully monitored experiments are rare, especially for bananas and BBTV, which is why we used data from two distant countries for model validation. This approach not only tests the robustness of parameterisation from small experiments to larger-scale dynamics but also demonstrates the ability to infer epidemic behavior across three spatially separated countries within sub-Saharan Africa.

Several limitations should be noted. First, the leaf emergence rate (LER) curve used for parameterisation was based on field data from sub-tropical Australia (Allen et al., 1988); changes in the LER curve could shift predicted optimal surveillance timing. Second, we assumed that all infected plants eventually express symptoms and that inspections would therefore achieve 100% detection. In practice, early disease symptoms are difficult to detect without training, so detection probabilities would vary depending on surveyor expertise. Future work should explicitly incorporate detection probability to reflect real-world surveillance conditions more accurately and to improve evaluation of management strategies.

In conclusion, our study demonstrates that small field experiments can provide sufficient data to parameterise epidemiological models, enabling both mechanistic understanding of infection dynamics and informed preparation for future outbreaks. By linking detailed modeling with practical field-level considerations, these approaches can support improved surveillance and targeted management of BBTV in banana-growing regions.

## Data Availability

Analysis code used in this study can be accessed at the following URL: https://github.com/rretkute/BBTVModelSmallFieldData.

